# Deep time course proteomics of SARS-CoV- and SARS-CoV-2-infected human lung epithelial cells (Calu-3) reveals strong induction of interferon-stimulated gene (ISG) expression by SARS-CoV-2 in contrast to SARS-CoV

**DOI:** 10.1101/2021.04.21.440783

**Authors:** Marica Grossegesse, Daniel Bourquain, Markus Neumann, Lars Schaade, Andreas Nitsche, Joerg Doellinger

**Affiliations:** Robert Koch Institute, Centre for Biological Threats and Special Pathogens: Highly Pathogenic Viruses (ZBS 1); Robert Koch Institute, Centre for Biological Threats and Special Pathogens; Robert Koch Institute, Centre for Biological Threats and Special Pathogens: Proteomics and Spectroscopy (ZBS 6)

**Keywords:** SARS-CoV-2, coronavirus, interferon response, interferon-stimulated gene (ISG), proteomics, data-independent-acquisition

## Abstract

SARS-CoV and SARS-CoV-2 infections are characterized by remarkable differences, including contagiosity and case fatality rate. The underlying mechanisms are not well understood, illustrating major knowledge gaps of coronavirus biology. In this study, protein expression of SARS-CoV- and SARS-CoV-2-infected human lung epithelial cell line Calu-3 was analysed using data-independent acquisition mass spectrometry (DIA-MS). This resulted in the so far most comprehensive map of infection-related proteome-wide expression changes in human cells covering the quantification of 7478 proteins across 4 time points. Most notably, the activation of interferon type-I response was observed, which surprisingly is absent in other recent proteome studies, but is known to occur in SARS-CoV-2-infected patients. The data reveal that SARS-CoV-2 triggers interferon-stimulated gene (ISG) expression much stronger than SARS-CoV, which reflects the already described differences in interferon sensitivity. Potentially, this may be caused by the enhanced expression of viral M protein of SARS-CoV in comparison to SARS-CoV-2, which is a known inhibitor of type I interferon expression. This study expands the knowledge on the host response to SARS-CoV-2 infections on a global scale using an infection model, which seems to be well suited to analyse innate immunity.

## Introduction

In late 2019, first cases of severe pneumonia of unknown origin were reported in Wuhan, China. Shortly afterwards a new coronavirus was discovered as the causative agent and named SARS-CoV-2 and the related disease COVID-19. The virus turned out to be highly contagious and caused a world-wide pandemic, which is still ongoing and has already led to the death of > 2,900,000 humans worldwide. Already in 2002, another coronavirus, SARS-CoV, was discovered in China which was related to a severe acute respiratory syndrome (SARS) and caused an outbreak with about 780 deaths (1). However, at this time the outbreak could be controlled probably due to the lower contagiosity of SARS-CoV compared to SARS-CoV-2 (2). SARS-CoV and SARS-CoV-2 share about 80 % of their genome sequence and protein homology ranges between 40 and 94% (3, 4). Although both viruses mainly lead to respiratory tract infections and can cause severe pneumonia, they are characterized by remarkable differences, including contagiosity and case fatality rate (5). As the respiratory tract is the first and main target of SARS-CoV and SARS-CoV-2 infections, it seems conclusive to use airway epithelia cells to study differences of these two viruses. However, no comparative proteomics study has been published using Calu-3 cells which is the only permissive lung cell line available for SARS-CoV and SARS-CoV-2 (6). Other human lung cells lines, like A549, are only susceptible to SARS-CoV-2 infection upon overexpression of the SARS-CoV receptor ACE2 (6) which was recently found to be an interferon-stimulated gene (ISG) (7). In the present study, we used data-independent acquisition mass spectrometry (DIA-MS) to analyse the protein expression in Calu-3 cells infected with SARS-CoV and SARS-CoV-2 over the time course of 24 hours. In total, 8391 proteins were identified, 7478 of which could be reliably quantified across the experiment. This results in a deep and comprehensive proteome map which reflects time-dependent protein expression changes during SARS-CoV and SARS-CoV-2 infections and provides deep insights into the virus-specific immunomodulation of human lung cells.

## Methods

### Cell culture and infection

Calu-3 cells (ATCC HTB-55) were cultivated in EMEM containing 10 % FCS, 2 mM L-Gln and non-essential amino acids. A total of 5×10^5^ cells per well were seeded in 6-well plates and incubated overnight at 37°C and 5% CO2 in a humified atmosphere. Medium was removed and cells were infected with SARS-CoV (strain Hong Kong) or SARS-CoV-2 (hCoV-19/Italy/INMI1-isl/2020 (National Institute for Infectious Diseases, Rome, Italy, GISAID Accession EPI_ISL_410545) at an MOI of 5. Mock samples were treated with medium only. After one hour post infection (p.i.) cells were washed with PBS and fresh medium was added. After 2, 6, 8, 10 and 24 h p.i. the medium was removed and stored at -80 °C. Cells were washed with PBS and prepared for proteomics as described below. For each time point and virus, triplicate samples were taken. Additionally, triplicate mock samples per time point were taken.

### Polymerase chain reaction (PCR)

The amount of SARS-CoV and SARS-CoV-2 RNA in the supernatant was analysed by qPCR at 2, 6, 8, 10 and 24 h p.i.. Supernatants were extracted using the QIAamp Viral RNA Mini Kit (Qiagen, Hilden, Germany) according to manufacturer’s recommendations and eluted in 60 µL of RNase-free water. Real-time PCR targeting the viral E gene was carried out as described by Michel et al. (under revision) using the primers and probe published by Corman et al.(8). Quantification of viral genome equivalents (GE) was done using the SARS-CoV-2 E gene WHO reference PCR standard.

### IRF-activity reporter assay

ACE2-A549-Dual™ cells were seeded into 96-well plates at 4×10^4^ cells per well and incubated overnight at 37°C and 5% CO2 in a humified atmosphere. Cells were infected with either SARS-CoV or SARS-CoV-2 at an MOI of 1.0. At 2 d p.i., interferon regulatory factor (IRF)-activity was assayed using QUANTI-Luc™ luminescence reagent (InvivoGen, San Diego, CA, USA) and an INFINITE 200 PRO microplate reader (Tecan, Männedorf, Switzerland).

#### Sample preparation for proteomics

Samples were prepared for proteomics using Sample Preparation by Easy Extraction and Digestion (SPEED) (9). At first, medium was removed and cells were washed using phosphate-buffered saline. Afterwards, 200 µL of trifluoroacetic acid (TFA) (Thermo Fisher Scientific, Waltham, MA, USA) were added and cells were incubated at room temperature for 3 min. Samples were neutralized by transferring TFA to prepared reaction tubes containing 1.4 mL of 2M TrisBase. After adding Tris(2-carboxyethyl)phosphine (TCEP) to a final concentration of 10 mM and 2-Chloroacetamide (CAA) to a final concentration of 40 mM, samples were incubated at 95°C for 5 min. 200 µL of the resulting solutions were diluted 1:5 with water and subsequently digested for 20 h at 37°C using 1 µg of Trypsin Gold, Mass Spectrometry Grade (Promega, Fitchburg, WI, USA). Resulting peptides were desalted using 200 µL StageTips packed with three Empore™ SPE Disks C18 (3M Purification Inc., Lexington, USA) and concentrated using a vacuum concentrator (10, 11). Dried peptides were suspended in 20 µL of 0.1 % TFA and quantified by measuring the absorbance at 280 nm using an Implen NP80 spectrophotometer (Implen, Munich, Germany).

#### Liquid chromatography and mass spectrometry

Peptides were analysed on an EASY-nanoLC 1200 (Thermo Fisher Scientific, Bremen, Germany) coupled online to a Q Exactive™ HF mass spectrometer (Thermo Fisher Scientific). 1 µg of peptides were loaded on a μPAC™ trapping column (PharmaFluidics, Ghent, Belgium) at a flow rate of 2 µL/min for 6 min and were subsequently separated on a 200 cm μPAC™ column (PharmaFluidics) using a stepped 160 min gradient of 80 % acetonitrile (solvent B) in 0.1 % formic acid (solvent A) at 300 nL/min flow rate: 3–10 % B in 22 min, 10–33 % B in 95 min, 33–49 % B in 23 min, 49–80 % B in 10 min and 80 % B for 10 min. Column temperature was kept at 50°C using a butterfly heater (Phoenix S&T, Chester, PA, USA). The Q Exactive™ HF was operated in a data-independent (DIA) manner in the m/z range of 350–1,150. Full scan spectra were recorded with a resolution of 120,000 using an automatic gain control (AGC) target value of 3 × 10^6^ with a maximum injection time of 100 ms. The Full scans were followed by 84 DIA scans of dynamic window widths using an overlap of 0.5 Th (Table.S1). For the correction of predicted peptide spectral libraries, a pooled sample was measured using gas-phase separation (8 × 100 Th) with 25 × 4 Th windows in each fraction using a shift of 2 Th for subsequent cycles. Window placement was optimised using Skyline (Version 4.2.0) (11). DIA spectra were recorded at a resolution of 30,000 using an AGC target value of 3 × 10^6^ with a maximum injection time of 55 ms and a first fixed mass of 200 Th. Normalized collision energy (NCE) was set to 25 % and default charge state was set to 3. Peptides were ionized using electrospray with a stainless steel emitter, I.D. 30 µm (Proxeon, Odense, Denmark) at a spray voltage of 2.0 kV and a heated capillary temperature of 275°C.

### Data analysis

Protein sequences of *Homo sapiens* (UP000005640, 95,915 sequences, downloaded 23/5/19), SARS-CoV (UP000000354, 15 sequences, downloaded 21/9/20) and SARS-CoV-2 (UP000464024, 14 sequences, downloaded 21/9/20) were obtained from UniProt (12). A combined spectral library was predicted for all possible peptides with strict trypsin specificity (KR not P) in the m/z range of 350–1,150 with charge states of 2–4 and allowing up to one missed cleavage site using Prosit (13).

Input files for library prediction were generated using EncyclopeDIA (Version 0.9.5) (14). The *in-silico* library was corrected using the data of the gas-phase fractionated pooled sample in DIA-NN (Version 1.7.10) (15). Mass tolerances were set to 10 ppm for MS1 and 20 ppm for MS^2^ spectra, and the “unrelated run” option was enabled with the false discovery rate being set to 0.01. The single-run data were analysed using the corrected library with fixed mass tolerances of 10 ppm for MS1 and 20 ppm for MS^2^ spectra with enabled “RT profiling” using the “robust LC (high accuracy)” quantification strategy. The false discovery rate was set to 0.01 for precursor identifications and proteins were grouped according to their respective genes. The resulting identification file was filtered using R (Version 3.6) in order to keep only proteotypic peptides and proteins with protein q-values < 0.01. Visualization and further analysis were done in Perseus (Version 1.6.5) (16). Relative protein quantification was done based on log (2)-transformed and Z-score normalized “MaxLFQ” intensities. Proteins which were not quantified in at least 2/3rd of all samples were removed, and remaining missing values were replaced from a normal distribution (width 0.3, down shift 1.8). Significant protein expression differences between samples were identified using an ANOVA test with a permutation-based FDR of 0.05 (250 randomizations, s0 = 1). Afterwards a post-hoc test was applied to detect significant sample pairs using an FDR of 0.05. Gene ontology enrichment of differentially expressed proteins was analysed using the ClueGO app (Version 2.5.7) implemented in Cytoscape (Version 3.8.2) with a Bonferroni-adjusted p-value threshold of 0.05 (12, 17, 18).

## Results

Proteome analysis of SARS-CoV- and SARS-CoV-2-infected human lung epithelial cell line Calu-3 was conducted at 2, 6, 10 and 24 h p.i. including time-matched mock controls. Samples were prepared as biological triplicates and analysed using single-shot DIA-based proteomics with an optimized workflow for deep and accurate protein profiling (19). In total, 8391 proteins were identified in a 3 h gradient of which 7478 proteins were consistently quantified and used for further analysis (Pearson correlation > 0.98, median coefficient of variation between 0.048–0.062 within each triplicate, data completeness 98.3 %). Viral replication was verified by qPCR of the cell culture supernatants. The number of viral genome copies started to increase 6 h p.i. and no difference among SARS-CoV and SARS-CoV-2 was observed at any time point (**Figure 1**). This is consistent with the expression of viral proteins, which was detectable from 6 h p.i. as well. The majority of viral proteins including nucleoprotein, spike glycoprotein, ORF3a, ORF7a and ORF9a are expressed in equal amounts upon infection with both viruses. The only exception is the membrane protein (M) whose expression is enhanced in SARS-CoV-infected cells compared to SARS-CoV-2-infected ones (**Figure 1**).

**Figure 1.**
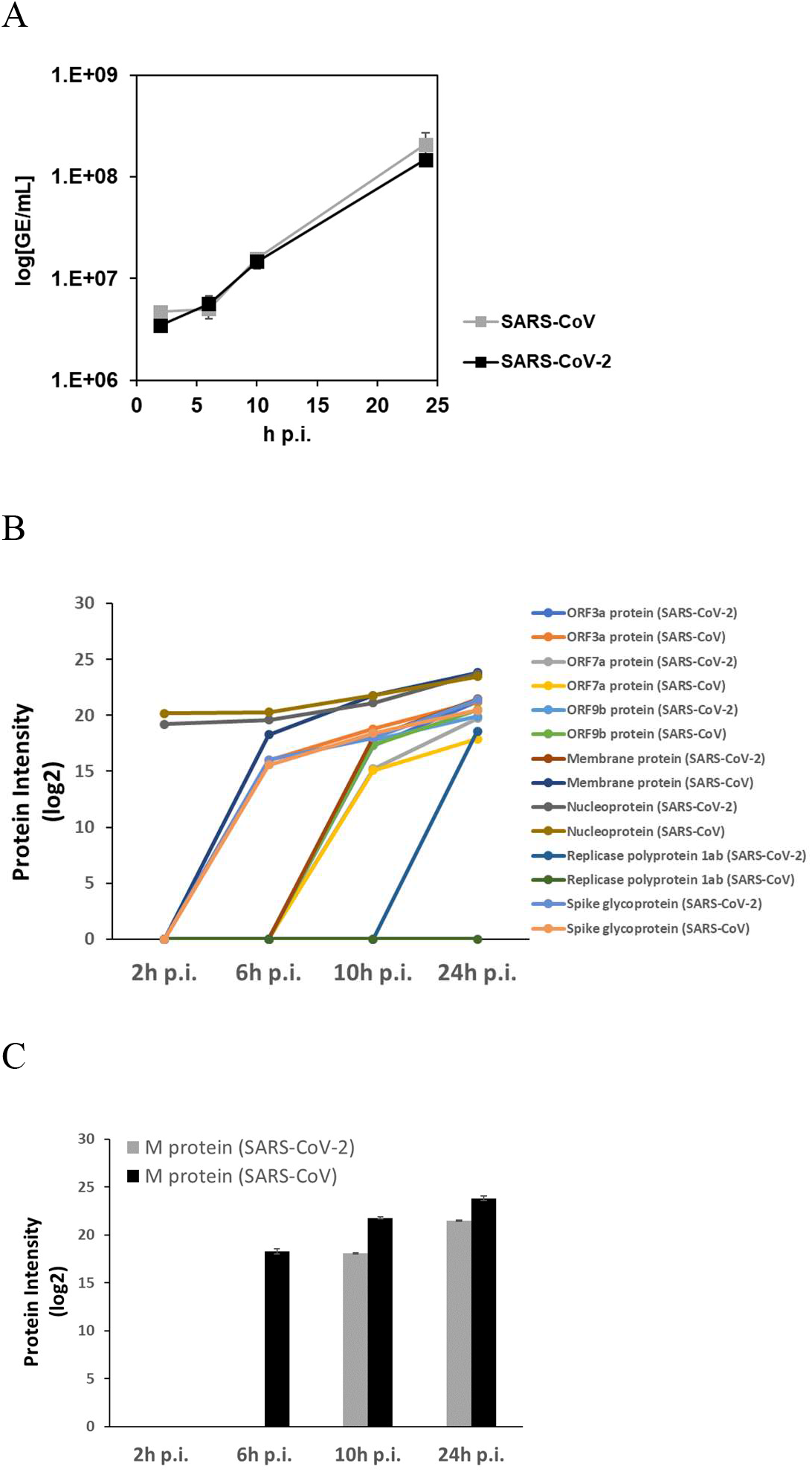
Viral protein expression and quantification of virus in the supernatant. Calu-3 cells were infected with SARS-CoV, SARS-CoV-2 or mock infected. After 2, 6, 10 and 24 h post infection (p.i.) the virus was quantified in the supernatant by qPCR (A). Protein expression in infected cells was analysed by data-independent acquisition (DIA) mass spectrometry. Intensities of viral proteins in infected Calu-3 cells are shown in (B). Expression of viral M protein is shown in (C).

The expression of 2642 human proteins differed significantly between the sample groups (ANOVA, FDR = 0.05), which was reduced to 261 proteins using a post-hoc test (FDR = 0.05) when only proteins with at least one significant pairwise difference in an infected cell with its time-matched mock control were kept. This large reduction underlines the need for time-matched mock controls in viral proteomics as long incubation times themselves can already lead to large alterations of the cellular proteome. The remaining infection-related proteins were grouped using hierarchical clustering according to their expression profiles, and the respective main clusters were analysed for enriched gene ontology terms using ClueGO (**Figure 2**). Out of the five clusters two clusters (up-regulated 2 h p.i. and down-regulated 6 h p.i.) revealed no significantly enriched GO terms but among others contained several proteins related to immune response such as OAS1, INAVA and NFKBIB. Another cluster consisting of proteins with virus-specific time-course-dependent upregulation was found to be related to mitochondrial translation (adjusted p-value: 2.5 * 10^−4^, MRPL17, MRPL27, MRPL47, MRPL50 and MRPS7). The other two main clusters included upregulated proteins 24 h p.i. and are related to either the regulation of complement activation (adjusted p-value: 7.9 * 10^−3^, C3 and C5) or interferon alpha/beta signalling (adjusted p-value: 7.8 * 10^−20^, e.g. MX1, MX2, DDX58, STAT1, OAS2, OAS3 and IFIT3).

**Figure 2.**
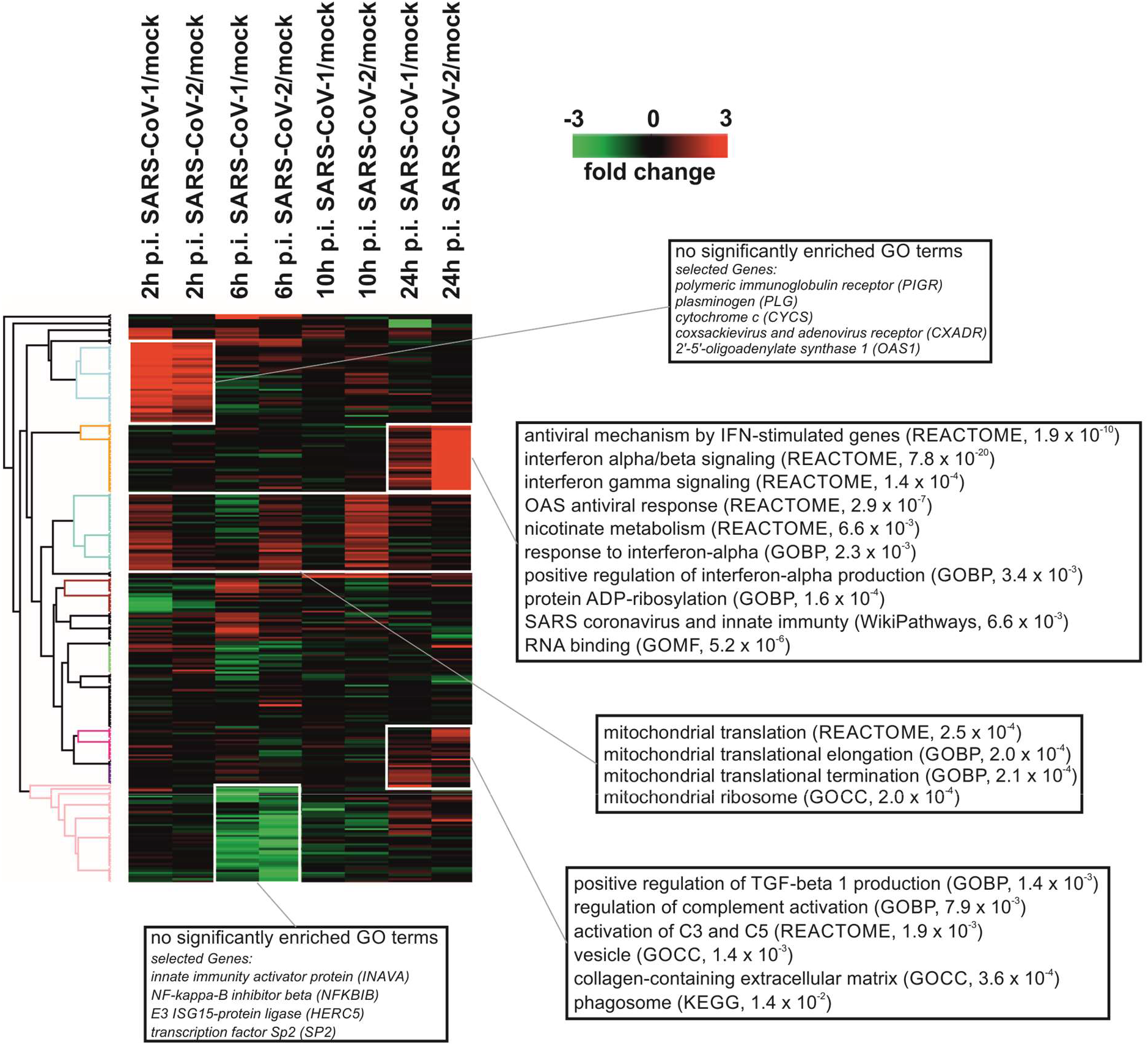
Infection-related alterations in the host proteome. Infection of Calu-3 cells with SARS-CoV and SARS-CoV-2 altered the abundance of 261 human proteins in comparison to time-matched mock controls. The heatmap depicts those proteins represented by their log2-transformed intensities using hierarchical clustering. Selected GO terms resulting from an enrichment analysis using ClueGO are denoted for the five main clusters. Complete results of the GO analysis can be found in the supplementary information.

Strikingly, the main difference between SARS-CoV- and SARS-CoV-2-infected cells was observed for proteins derived from interferon-stimulated genes (ISG), whose expression is enhanced in SARS-CoV-2-infected cells in comparison to SARS-CoV infection. This was confirmed by higher IFN induction triggered by SARS-CoV-2 in ACE2-A549 reporter cells compared to no detectable IFN-regulatory factor activity upon infection with SARS-CoV (**SI Figure 1**). As the type I interferon response is the most important one of the innate immune system to RNA viruses, we compared the expression data of related proteins from this study with other major proteome studies of SARS-CoV-2-infected human cells. For this purpose, all identified proteins annotated with the GO term “type I interferon signalling pathway” (GO:0060337) were extracted from the data of Stukalov et al. (https://covinet.innatelab.org, A549-ACE2 cells, MOI = 2) and Bojkova et al. (Caco-2 cells, MOI = 1), matched and clustered according to their expression profiles (**Figure 3**) (20, 21). The resulting heatmap revealed that the activation of the type I interferon response is completely absent in the other studies. However, it has to be noted that the coverage of this pathway differs strongly among the studies. Most of the ISGs with expression changes induced by infection are exclusively detected in the present study, which reflects the fact that the total number of quantified proteins was the largest among the studies as well. Furthermore, an interaction network of all infection-related proteins from this study was constructed using STRING ((22), https://string-db.org/) (**Figure 4**). The network revealed high connectivity among proteins related to either innate immunity (mainly interferon type I signalling), exocytosis, including proteins related to platelet degranulation (adjusted p-value: 0.01, e.g. FGB, FGG, FN1, PLG and PSAP) or mitochondria-associated proteins including many members of the ribonucleoprotein complex related to mtDNA expression.

**Figure 3.**
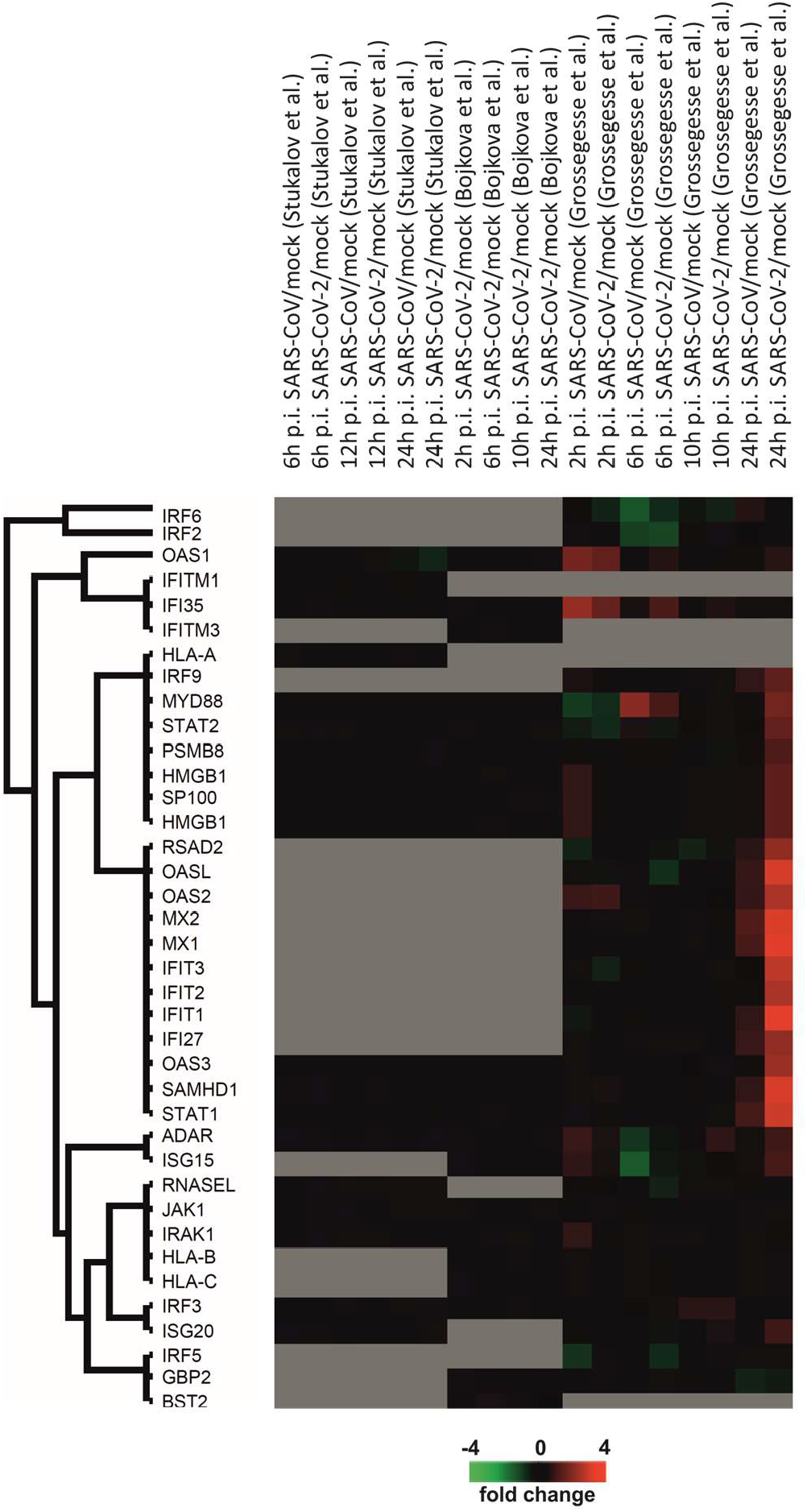
Meta-analysis of proteins associated with type I IFN signalling pathway. Expression data of all identified proteins associated with type I interferon signalling pathway (GO:0060337) were extracted from proteome studies of SARS-CoV-2-infected human cell lines done by Stukalov et al., Bojkova et al. and Grossegesse et al. and summarized in a heatmap representing log2-transformed intensity values. Missing values are grey.

**Figure 4.**
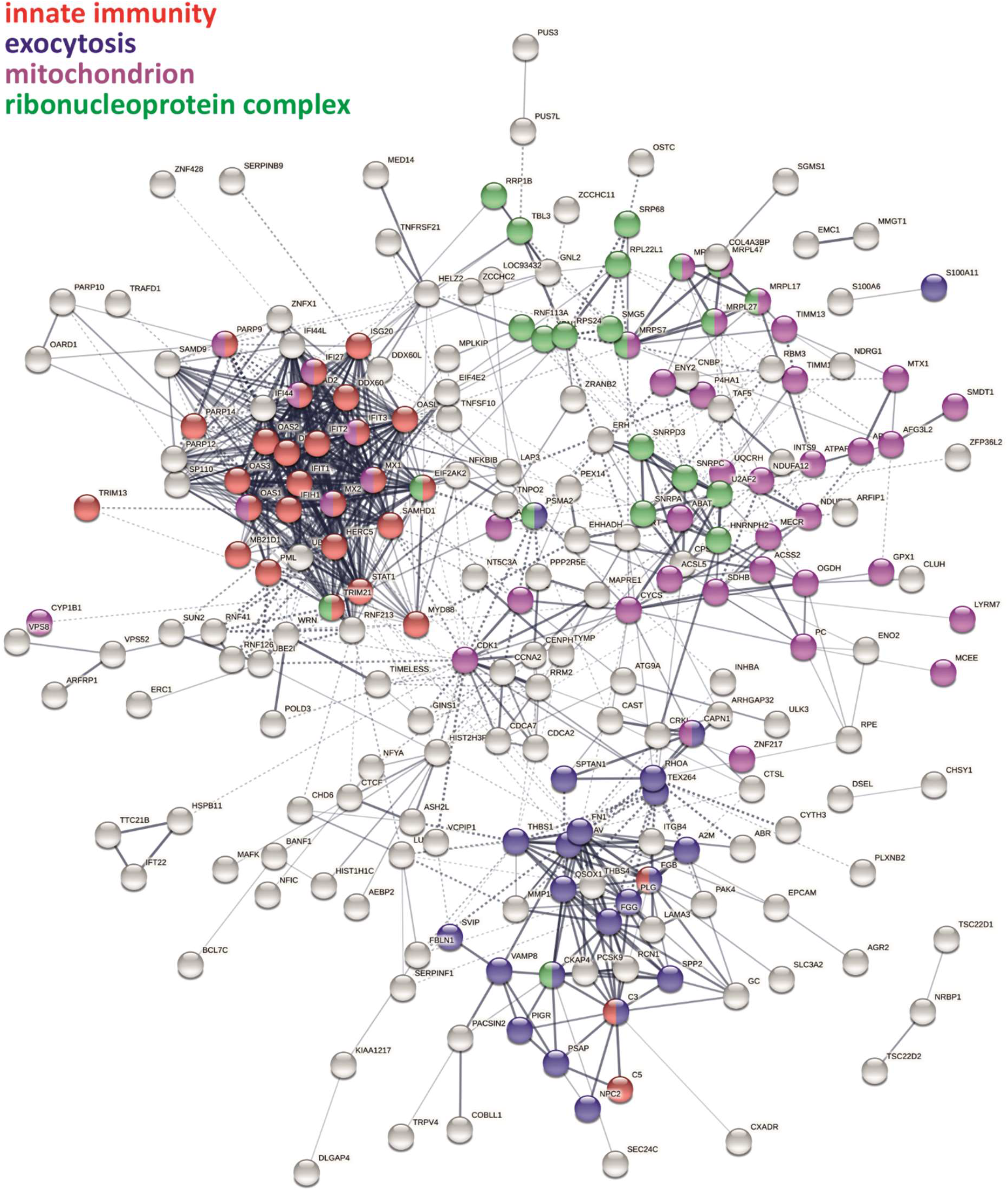
Protein interaction network of infection-related human proteins. The interaction network of all human proteins (N = 261) in Calu-3 cells affected by SARS-CoV and SARS-CoV-2 infections was constructed using StringDB.

## Discussion

Innate immunity is the host’s first line of defence to fight infections. The most important mechanism to combat replication of RNA viruses is the interferon response. It is based on the recognition of pathogen-associated molecular patterns (PAMP), especially double-stranded RNA (dsRNA), which in the end results in the secretion of type-I interferons which in turn induce the expression of interferon-stimulated genes (ISG) including multiple antiviral proteins (23). Recently, it was shown that SARS-CoV-2 is more sensitive to both IFN-α and IFN-β treatment in cultured cells than SARS-CoV (24-27), which could favour a positive outcome of several clinical trials evaluating type-I IFNs as a possible treatment for COVID-19 (28). Clinical data from SARS-CoV-2-infected patients report low or absent levels of IFN-I in serum but induction of ISG expression (3, 29). It was further demonstrated that SARS-CoV-2 induces types I, II or III interferons in infected human lung tissues in contrast to SARS-CoV (30). However, the mechanism behind the varying IFN sensitivity of closely-related SARS-CoV and SARS-CoV-2 is elusive. In general, proteomics should be well suited to uncover the modulation of the type-I interferon response by SARS coronaviruses.

The experiments in the present study resulted in the so far most comprehensive map of infection-related proteome expression changes in SARS-CoV- and SARS-CoV-2-infected cells covering ∼ 7400 proteins across 4 time points. Expression of 261 proteins changed during the course of infection, which cluster into 5 main groups. One of those clusters reveals a strong induction of ISG expression 24 h p.i. in SARS-CoV-2-infected cells. Strikingly, this induction was observed at a much lower level in SARS-CoV-infected cells, which could reflect the varying IFN sensitivity. Among those ISG proteins is e.g. Mx1 which is known for its antiviral activity against a wide range of viruses. It was shown before that Mx1 expression is increased in SARS-CoV-2-infected patients and correlates well with viral load (31). Furthermore, it was demonstrated that ISG expression is induced in SARS-CoV-2-infected patients in general and that the increase of ISG expression, including Mx1, has a negative correlation with disease severity (29). Surprisingly, these findings are not reflected in the current literature of large-scale proteome analysis of infected human cells (20, 21). The absence of an enhanced ISGs expression in other proteome studies can result from incomplete proteome coverage or from different experimental conditions, e.g. different cell lines and MOIs. It was shown before that ISGs and IFN can be detected upon infection of A549-ACE2 and Calu-3 cells with SARS-CoV-2 and that higher MOIs favour interferon induction (32, 33). However, this was surprisingly not detected in the study of Stukalov et al. To shed light on this discrepancy, we performed a meta-analysis of type-I interferon related proteins by comparing data from this study to the studies of Bojkova et al. and Stukalov et al. (20, 21).

Interestingly, most of the strongly affected ISGs, including Mx1, Mx2, IFIT1, IFIT2, IFIT3, OASL and OASL2, were not identified in the previous studies. The low coverage of this pathway could explain at least partially the discrepancy. It must also be noted that the influence of ACE2 overexpression, which was used by Stukalov et al. to turn A549 into a permissive cell line, on the immune response is unknown, and recently it has been shown that ACE2 is an ISG itself (7). This meta-analysis demonstrates that proteome coverage is still a limitation which impedes intra-study cross-comparisons due to missing values.

Recently, it was proposed that SARS-CoV-2 ORF6 interferes less efficiently with human interferon induction and interferon signalling than SARS-CoV ORF6, which could explain the virus-specific induction of ISG expression and the varying interferon sensitivity (34). The proteome data from this study point towards an additional mechanism. The expression of viral proteins was highly similar between SARS-CoV and SARS-CoV-2 except for the M protein whose expression is enhanced in SARS-CoV. This protein is a component of the viral envelope but its functions beyond are not well characterized. It is known that the homologous M proteins of MERS and SARS-CoV inhibit type I interferon expression (35, 36). Recently, it was discovered that overexpression of the M protein from SARS-CoV-2 in human cells inhibits the production of type I and III IFNs induced by dsRNA-sensing via direct interaction with RIG-I (DDX58) and reduces the induction of ISGs after Sendai virus (SEV) infection and poly (I:C) transfection (33, 37). Additionally, it was shown that the M protein of SARS-CoV inhibits the formation of TRAF3·TANK·TBK1/IKKϵ complex, resulting in the inhibition of IFN transcription (35). We therefore hypothesize that the enhanced expression of the M protein of SARS-CoV reduces the induction of ISG expression in infected cells in comparison to SARS-CoV-2 and so contributes to the varying IFN sensitivity of both viruses. The gene expression of coronaviruses is controlled both on transcriptional and translational level (38). When comparing the core regulating elements of the M gene of SARS-CoV and SARS-CoV-2, it can be noted that both viruses have identical transcription regulatory sequences but quite diverse sequences around the translation initiation site, leading to the hypothesis of a different translational regulation (39). However, it should be noted that also sequence differences in the M protein of both viruses could lead to differences in the interferon-antagonizing capacity which is not known so far (**SI Figure 2**).

Summarized this study presents the so far most comprehensive comparative quantitative proteomics data set of SARS-CoV- and SARS-CoV-2-infected Calu-3 cells which are the only permissive human lung cell line for SARS-CoV-2 (6). By showing a diverse regulation of ISG expression upon infection, we conclude that Calu-3 cells present a good model system for studying differences in IFN sensitivity of SARS-CoV and SARS-CoV-2.

---

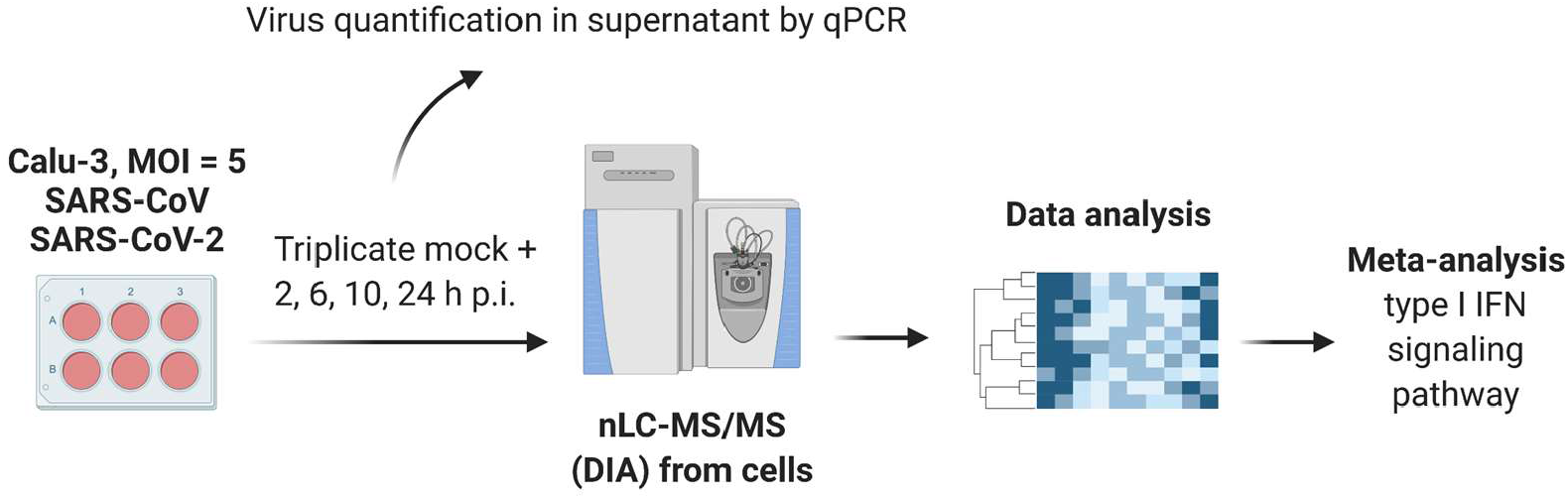

Graphical Abstract

**SI Figure 1.**
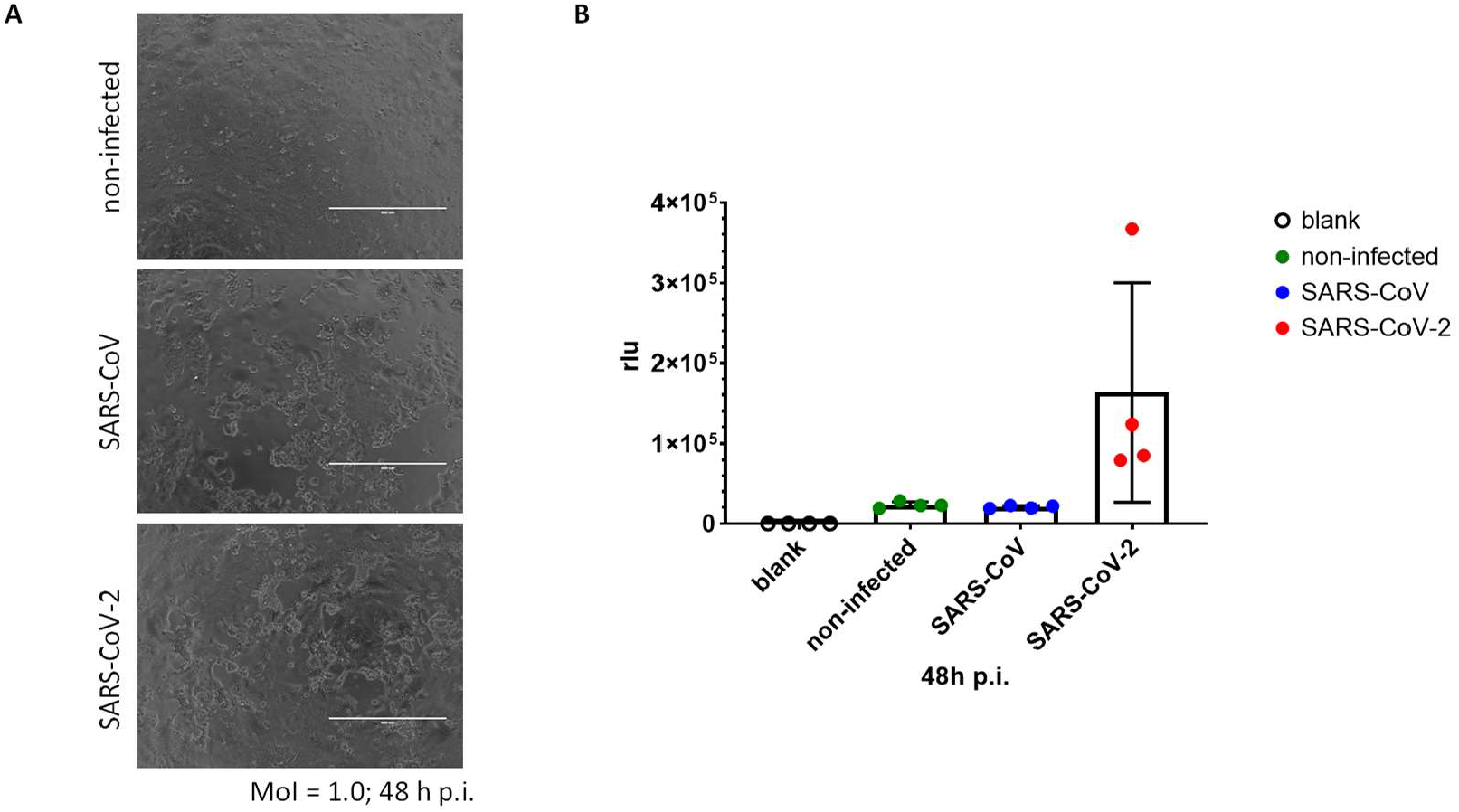
(A) Cytopathic effect and (B) IRF activity 48 h post SARS-CoV and SARS-CoV-2 infection of A549-Dual-ACE2 cells (MOI = 1.0).

**SI Figure 2:**
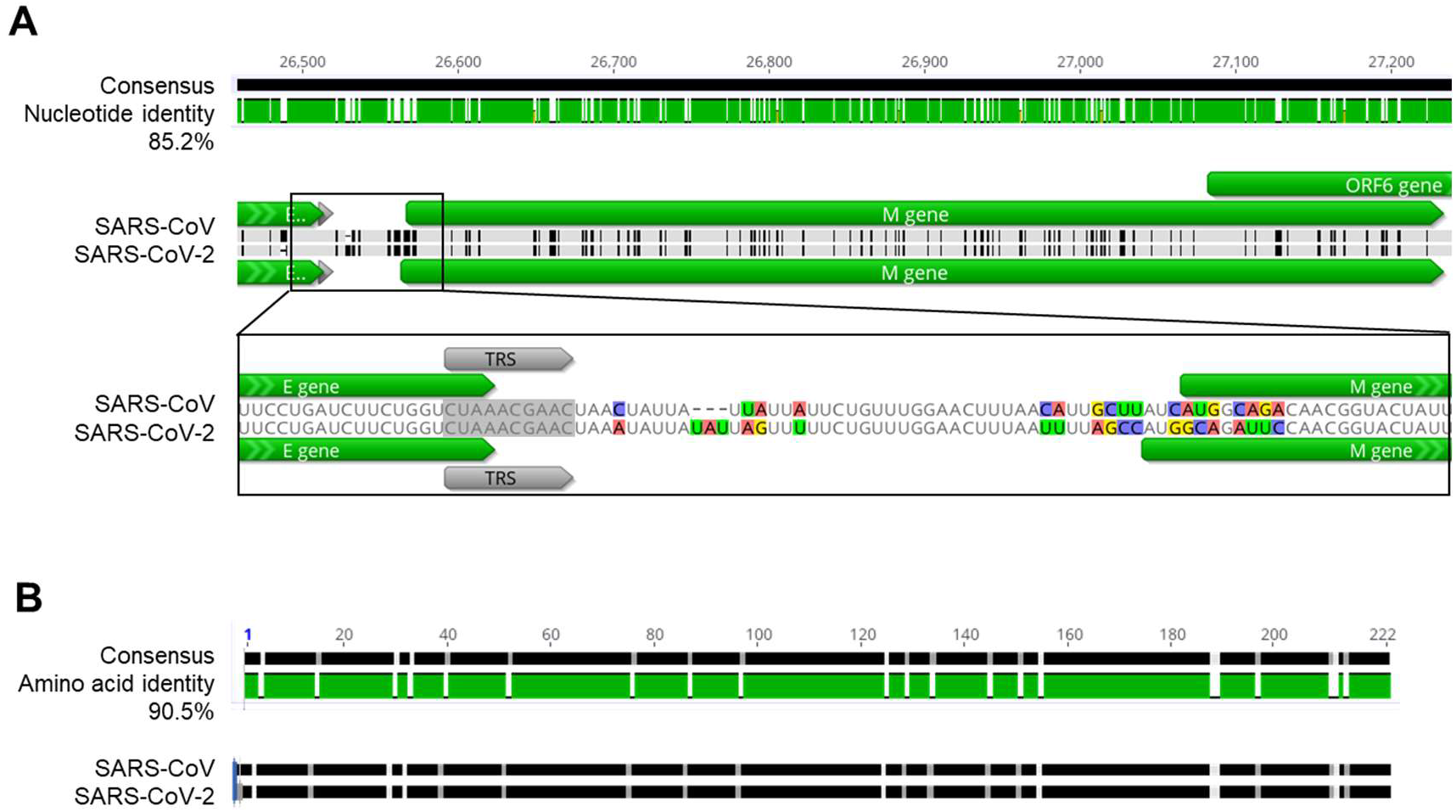
Sequence comparison of regulatory elements of the M protein of SARS-CoV and SARS-CoV-2. (A) Comparison of nucleotide sequences. TRS: transcription regulatory sequence according to Wu et al. Sequences were derived from NCBI: NC_004718 (SARS-CoV) and NC_045512 (SARS-CoV-2). (B) Comparison of amino acid sequences. Sequences were derived from UniProt: sp|P59596|VME1_SARS (SARS-CoV) and sp|P0DTC5|VME1_SARS2 (SARS-CoV-2).

## Acknowledgements

The authors would like to thank Clemens Bodenstein, Bianca Hube, Melanie Hoffmeister and Stefanie Schürer for supporting the infection experiments and qPCR and Ursula Erikli for copy-editing.

## Access to proteomics data

The mass spectrometry proteomics data have been deposited to the ProteomeXchange Consortium (http://proteomecentral.proteomexchange.org) via the PRIDE partner repository with the dataset identifiers PXD024883.

